# Computational Model to Quantify the Growth of Antibiotic Resistant Bacteria in Wastewater

**DOI:** 10.1101/2020.10.09.333575

**Authors:** Indorica Sutradhar, Carly Ching, Darash Desai, Mark Suprenant, Emma Briars, Zachary Heins, Ahmad S. Khalil, Muhammad H. Zaman

## Abstract

Although wastewater and sewage systems are known to be significant reservoirs of antibiotic resistant bacterial populations and periodic outbreaks of drug resistant infection, there is little quantitative understanding of the drivers behind resistant population growth in these settings. In order to fill this gap in quantitative understanding of the development of antibiotic resistant infections in wastewater, we have developed a mathematical model synthesizing many known drivers of antibiotic resistance in these settings to help predict the growth of resistant populations in different environmental scenarios. A number of these drivers of drug resistant infection outbreak including antibiotic residue concentration, antibiotic interaction, chromosomal mutation and horizontal gene transfer, have not previously been integrated into a single computational model. We validated the outputs of the model with quantitative studies conducted on the eVOLVER continuous culture platform. Our integrated model shows that low levels of antibiotic residues present in wastewater can lead to increased development of resistant populations, and the dominant mechanism of resistance acquisition in these populations is horizontal gene transfer rather than acquisition of chromosomal mutations. Additionally, we found that synergistic antibiotic interactions lead to increased resistant population growth. These findings, consistent with recent experimental and field studies, provide new quantitative knowledge on the evolution of antibiotic resistant bacterial reservoirs, and the model developed herein can be adapted for use as a prediction tool in public health policy making, particularly in low income settings where water sanitation issues remain widespread and disease outbreaks continue to undermine public health efforts.

**Importance:** The rate at which antimicrobial resistance (AMR) has developed and spread throughout the world has increased in recent years, and according to the Review on Antimicrobial Resistance in 2014 it is suggested that the current rate will lead several million people AMR-related deaths by 2050^25^. One major reservoir of resistant bacterial populations that has been linked to outbreaks of drug resistant bacterial infections, but is not well understood, is in wastewater settings, where antibiotic pollution is often present. Using ordinary differential equations incorporating several known drivers of resistance in wastewater, we find that interactions between antibiotic residues and horizontal gene transfer significantly affect the growth of resistant bacterial reservoirs.

## Introduction

Wastewater and sewage systems are a major reservoir of resistant bacterial populations due to the collection of antibiotic waste from humans and animals, inappropriate drug disposal, and effluent from drug manufacturers, hospitals and agricultural/veterinary settings (1). In some cases, environmental concentrations can reach, or even exceed, minimal inhibitory concentrations of certain antibiotics (2). This problem is particularly pertinent in low- and middle-income countries (LMICs) where cases of antibiotic resistant infections have been rising and 70% of sewage produced is estimated to enter the environment untreated (1). For example, in 2016, an outbreak of extensively drug resistant (XDR) typhoid cases emerged in in the southern Sindh province of Pakistan, and geospatial mapping revealed that the XDR *Salmonella* Typhi infections spread around sewage lines in the city of Hyderabad (3). Such outbreaks put the health and safety of surrounding populations at risk and, due to the communicable nature of these infections, increase risk for populations beyond the immediate vicinity of wastewater and sewage lines. Antibiotic pollution in wastewater and sewage systems results in a complex environment with many interacting antibiotic residues and this pollution is a major driver of the antibiotic resistance that is leading to outbreaks in surrounding communities (4, 5). One reason for this is that selective pressure from antibiotics in the environment is known to promote chromosomal resistance mutations (2, 6, 7). Furthermore, the interactions between different antibiotic residues may also affect resistance due to the synergistic and antagonistic effects changing the selective pressure on the bacteria (8). Additionally, the concentrations of antibiotic residues and disinfectants in sewage may be able to support and promote horizontal transfer of resistance genes among bacteria (9-12). These mobile resistance genes have previously been measured at high levels in wastewater and sewage (2). Though AMR and its drivers have been studied in sewage and wastewater settings to some extent (2, 12, 13), one of the largest gaps in understanding the emergence of AMR within a sewage environment is the limited quantitative understanding of the effects and interactions of the many biological and environmental mechanisms at work (14). Quantitative understanding of resistant population growth in wastewater settings is critical in order to predict the development of large resistant populations in wastewater that may pose a risk to local populations of resistant infection outbreak. Furthermore, quantitative tools for understanding of resistance can be used to develop and model strategies for preventing the emergence of large resistant populations and therefore influence policy decisions. Therefore, there is a critical need for developing quantitative methods of probing AMR in wastewater environments. Our study is a step in filling this gap in knowledge.

Mathematical modelling has been an important tool in quantitatively studying AMR development at both an epidemiologic and mechanistic level (6, 8, 12, 15-17). Epidemiological studies have included approaches that investigate bacterial population dynamics in a number of biological contexts including biofilms (15). At the population level, much of AMR modeling deals with disease states in which resistance patterns are long established, with significant gaps in the study of resistance emergence in new pathogens and the role of environmental factors in the development of AMR (16). At the mechanistic level, modelling has been used as a tool to understand the individual roles of both mutational and plasmid-mediated resistances (8, 12). Models based on resistance developed from chromosomal mutations have been used to study the role of synergistic and antagonistic antibiotic interactions on the emergence of resistant populations, finding that increased synergy increases the likelihood of resistance acquisition (8). Additionally, studies using modelling as a tool to understand horizontal gene transfer (HGT) as a driver of AMR have found HGT, specifically through the mechanism of conjugation, to be a significant mode of resistance acquisition for *Escherichia coli* in settings including agricultural slurry (12). However, much of the work done recently has been focused on fitting data in the absence of antibiotic exposure (17). Understanding how bacteria evolve upon antibiotic exposure is important to develop strategies that prevent the emergence of resistance (6). Thus, there remains a need for quantitative modeling of both chromosomal mutation acquisition and HGT under selective pressure from antibiotics to fully understand AMR development in settings such as sewage and wastewater, where antibiotic residues are often present. Here we present a model of emergence and growth of drug resistant bacteria in wastewater, integrating acquisition of resistance through both chromosomal mutations and HGT while also incorporating the effects of antibiotic interactions.

## Methods

Our mathematical model of the growth of antibiotic resistant bacterial populations in wastewater builds on prior studies (8, 12, 18) and extends it to incorporate a variety of critical, but overlooked, input factors specific to the bacterial species, antibiotics and environments of interest. The inputs can be broadly classified into bacterial parameters, environmental parameters and antibiotic parameters. Bacteria specific input factors include the growth rates of antibiotic susceptible and resistant strains, mutation rates, and rates of HGT. The antibiotic specific inputs, such as bactericidal activity and degree of synergy, allow for the study of the effects of drug quality and antibiotic pollution on the development of resistance. Additionally, environmental inputs, including physical fluid inflow and outflow rates and antibiotic residue concentration, allow for the modelling of resistance development in a variety of settings of interest. These input parameters can be used to model an output of resistant bacterial populations over time, thus allowing for the prediction of resistant population development (Figure 1).

**Figure 1.**
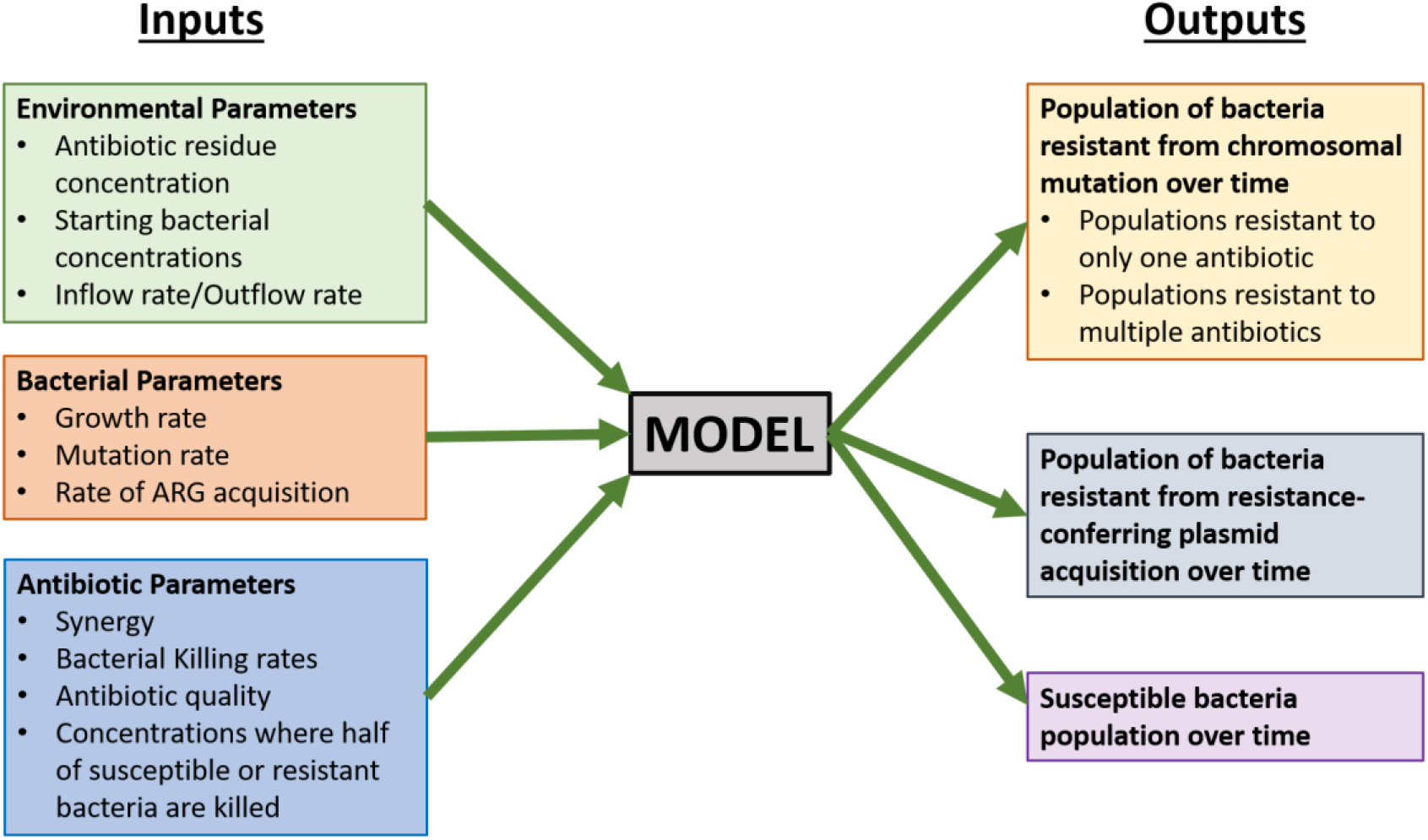
Inputs and outputs for preliminary model of antibiotic resistance development in a single bacterial species in a wastewater setting

The model consists of ordinary differential equations governing the concentration of two antibiotics (*C*_*1*_and C_*2*_) over time as well as equations modelling the susceptible population (*S)* and populations resistant to both antibiotics 1 and 2 from chromosomal mutation (*R*_*m*_) or from HGT, specifically a resistance-conferring plasmid (*R*_*p*_) over time (Eq set 1) (Table 1). Additionally, two populations of bacteria that are resistant through chromosomal mutation to each antibiotic individually are modelled (*R*_*1*_ and *R*_*2*_). Each antibiotic is modelled with terms for the antibiotic residue concentration in the environment (*E*) and the antibiotic clearance rate (*k*_*e*_). Each bacterial population is modelled with terms for growth rate (*α*), which is limited by the carrying capacity (*N*_*max*_). The bactericidal activities of the antibiotics are modelled by the term for killing rate in response to each antibiotic (*δ*_*max*,1_ *and δ*_*max*,2_). These killing terms are modified by the antibiotic concentration where the killing action is half its maximum value for either susceptibility 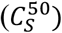 or resistance to each antibiotic 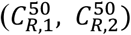. These killing rates and half max concentration terms are assumed to be equal for each antibiotic. Additionally, the killing rates are modified by a synergy term (*syn*) with *syn*<1 indicating antagonistic interaction and *syn*>1 indicating synergistic interaction. Also included in the model are terms for bacterial inflow (*g*_*s*_, *g*_*Rm*_, *g*_*R*1_, *g*_*R*2_, *g*_*Rp*_) and outflow (*k*_*T*_) rates (Table 2). These inflow and outflow rates are determined by the physical flow rates of the system of interest, and the outflow terms were validated experimentally as described below. As susceptible and resistant bacterial inflow and outflow rates for wastewater settings are not known, the model simulations assume bacterial inflow and outflow rates of zero. However, these parameters are still included due to their relevance for the modeling of systems such as wastewater where bacterial inflow and outflow will affect resistant population development and they may be quantified in future field studies. Chromosomal mutation is modelled through terms for mutation rates under antibiotic pressure (*m*) which are concentration dependent. Parameters governing mutation rate and antibiotic interaction are based on a previously developed model from Michel et al (8). HGT is modelled assuming the mechanism of plasmid conjugation and a HGT rate (*β*). HGT model terms and parameter values are adapted from a model of HGT in agricultural waste from Baker et al (12). Other model parameters are based on experimentally derived parameters of *E*. coli in piglet studies (18, 19). We note that the incorporation of all of these parameters, which have previously not been studied at once, into one single model can provide insights into their roles in the emergence and development of resistant populations. These equations were coded and solved in MATLAB (R2018a, Mathworks Inc).

**Table 1.**
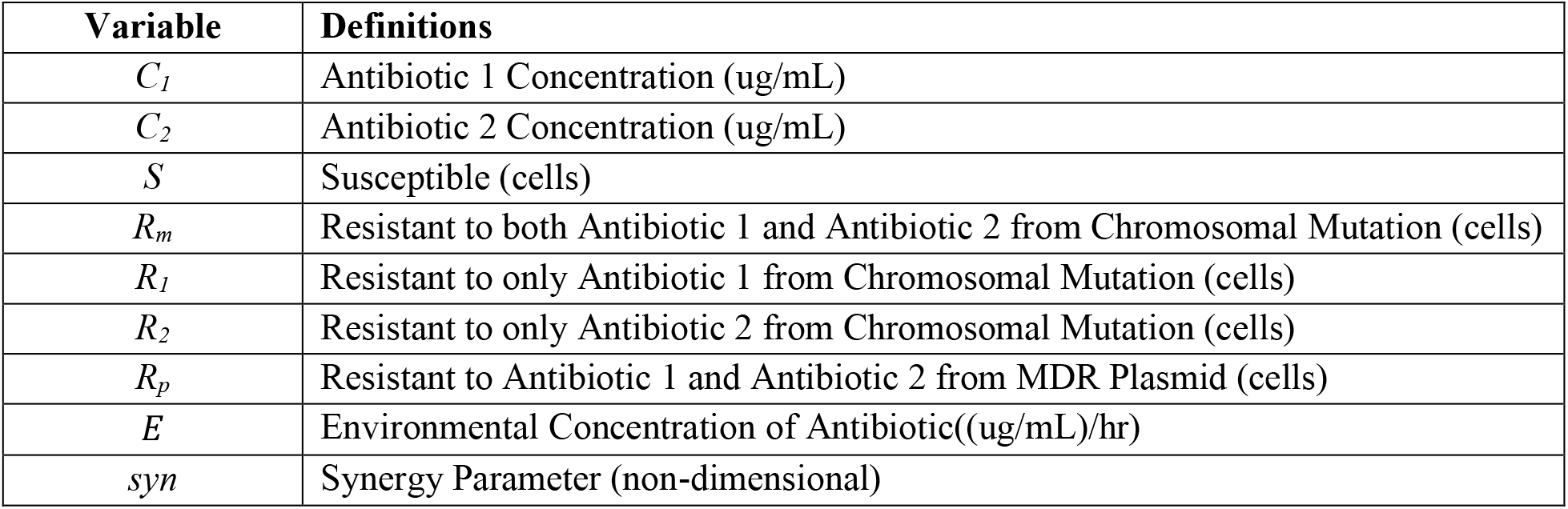
Model variables and definitions

**Table 2.**
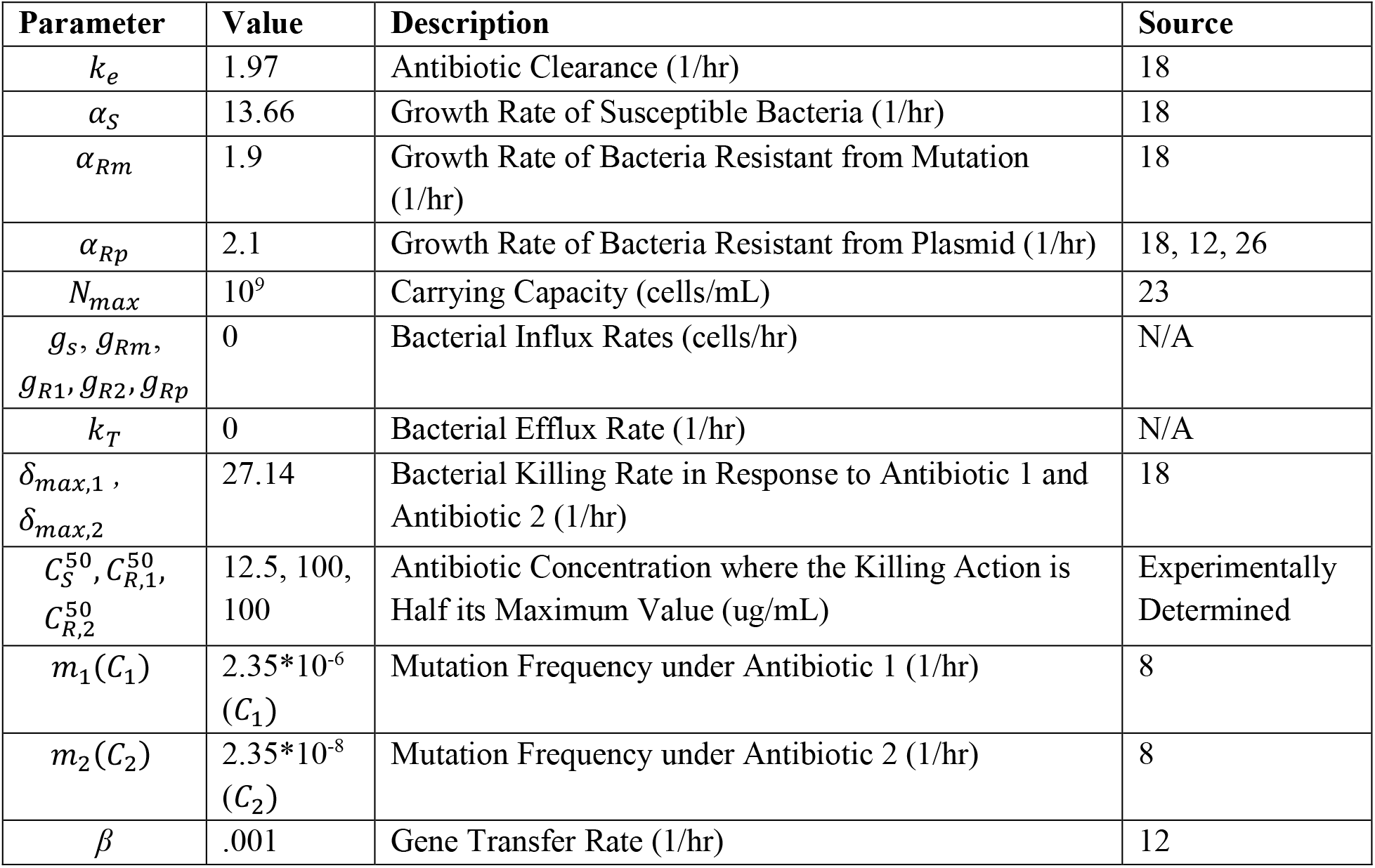
Model parameter values and descriptions

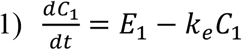

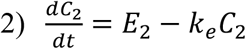

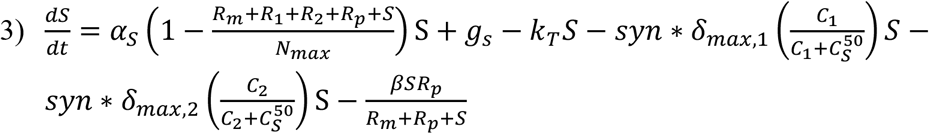

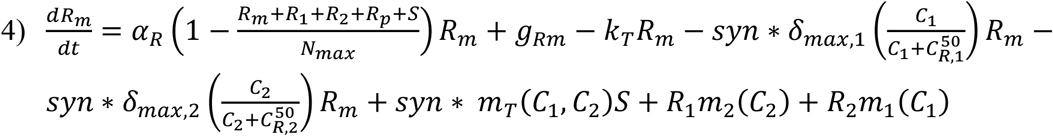

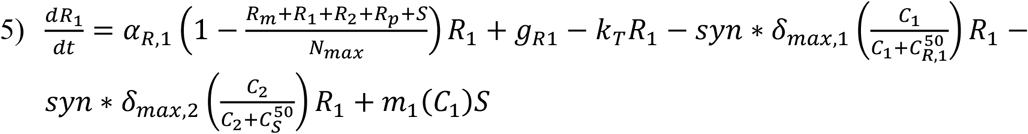

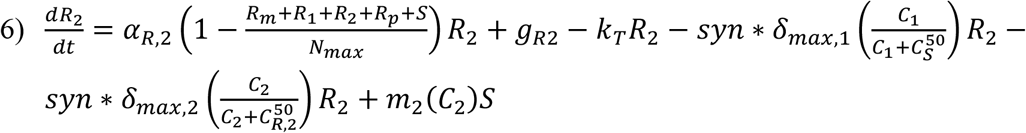

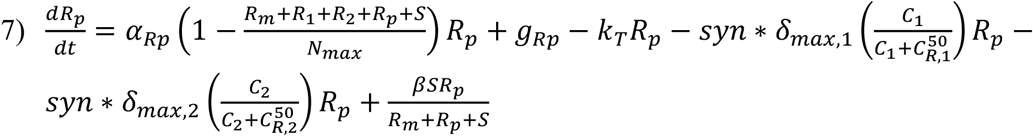

**Eq Set 1**. Sensitive and resistant populations under selective pressure from antimicrobial combination therapy including chromosomal mutation and HGT resistance mechanisms

Experimental validation of the model was done using the eVOLVER system (24). The experiment was initialized with inoculation of LB media with *E. coli* MG1655 in static conditions at 37°C. Then, inflow and outflow of Rifampicin-containing LB media at three concentrations (6, 8 and 12 mg/L) were started at a flow rate of 16 mL/hr. Following the beginning of inflow/outflow, each culture condition was sampled daily and the concentration of total bacteria was monitored through OD measurements taken through the eVOLVER system and resistance bacteria were calculated through plating on selective LB agar containing 200 mg/L Rifampicin. This Rifampicin concentration is eight-fold greater than the experimentally determined MIC for Rifampicin (25 mg/L).

## Results and Discussion

### Model Validation

In order to validate the developed model, experiments were conducted with a simplified experimental system including only one antibiotic (Rifampicin) and no plasmid-containing bacteria such that there was only chromosomal mutation as a mechanism for resistance. This experimental validation was done on the eVOLVER, a continuous culture platform which allows for automated, highly flexible and scalable microbial growth and lab evolution. eVOLVER allows for continuous flow conditions and independent, precise and multiparameter control of growth conditions such as temperature and flow rate (24). This allows us to precisely recreate the model parameters in vitro in a high throughput and repeatable manner. For these eVOLVER studies, we set certain inflow and outflow rates, initial bacterial populations, and antibiotic concentrations, as required as inputs for the computational model and then measure susceptible and resistant bacterial populations over the course of the continuous evolutions experiment. The outputs of the study can then be directly compared to the outputs of the model. The results of these studies are shown in Figure 2a-c. The experiments showed that at 6 mg/L and 8 mg/L Rifampicin (where the MIC of Rifampicin is 25 mg/L), the resistant population developed to a steady state around the same order of magnitude of the steady state population of susceptible bacterial. This steady state resistant population was larger for the 8 mg/L Rifampicin condition. At 12 mg/L, the experiments showed that the resistant bacteria dominated the population and the susceptible population dropped to zero. After the conclusion of the experiment, model equations and parameter values were adjusted to match the experimental behavior (Eq set 2) (Table 3). Notably, the mutation rates were increased by several orders of magnitude to match the rapid *in vitro* development of resistance. Additionally, the experimental results led to the addition of an equation for a theorized semi-resistant population with a lower level of resistance (MIC=100 mg/L) (S_1_). This was added to match the experimental finding that resistance can coexist at a steady state with the susceptible population. We theorized that the observed susceptible population also contained a subpopulation of semi-resistant cells which were not detected by the 8X MIC resistance cutoff rate of the experiment. These semi-resistant cells would allow for a sustained susceptible population in conditions with constant antibiotic pressure.

**Table 3.** Adjusted model parameter values and descriptions based on eVOLVER experimental conditions and results

| Parameter | Value | Description |
| --- | --- | --- |
| $k_e$ | 1.97 | Antibiotic Clearance (1/hr) |
| $\alpha_S$ | 13.66 | Growth Rate of Susceptible Bacteria (1/hr) |
| $\alpha_{R,1}$ | 4.99 | Growth Rate of Bacteria Resistant from Mutation (1/hr) |
| $\alpha_{S,1}$ | 8.3 | Growth Rate of Bacteria Semi-Resistant from Mutation (1/hr) |
| $N_{max}$ | $3 \times 10^8$ | Carrying Capacity (cells/mL) |
| $g_S, g_{R,1}, g_{S,1}$ | 0 | Bacterial Influx Rates (cells/hr) |
| $k_T$ | 0.8 | Bacterial Efflux Rate (1/hr) |
| $\delta_{max,1}, \delta_{max,2}$ | 27.14 | Bacterial Killing Rate in Response to Antibiotic 1 and Antibiotic 2 (1/hr) |
| $C_S^{50}, C_{S,1}^{50}, C_{R,1}^{50}$ | 12.5, 100, 200 | Antibiotic Concentration where the Killing Action is Half its Maximum Value (ug/mL) |
| $m_1(C_1)$ | $8.5 \times 10^{-3} (C_1)$ | Mutation Frequency of Full Resistance Mutation (1/hr) |
| $ms_1(C_2)$ | $8.5 \times 10^{-2} (C_1)$ | Mutation Frequency of Low Resistance Mutation (1/hr) |

**Figure 2.**
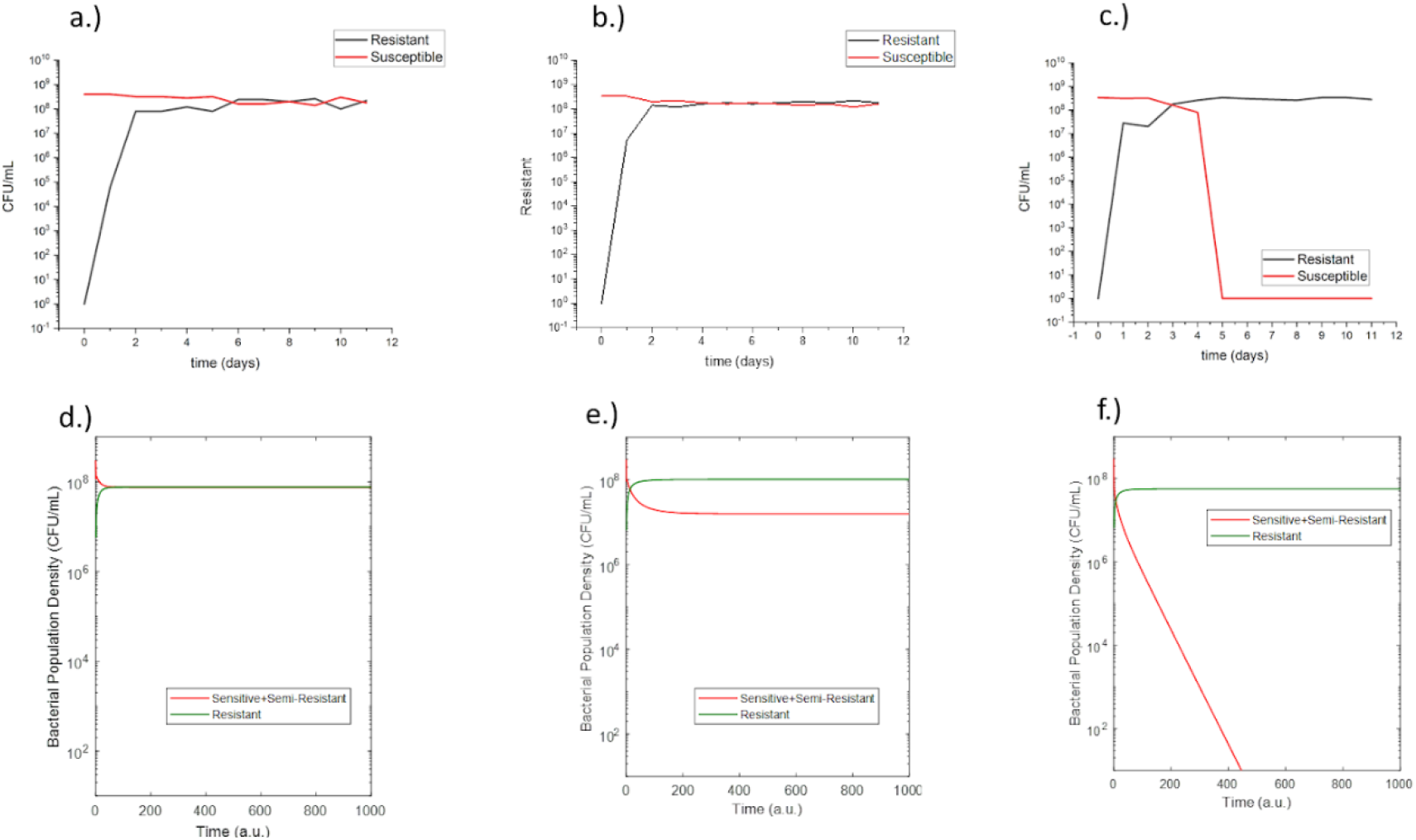
Measured susceptible and resistant populations from eVOLVER experiments of E. coli grown in continuous culture in media containing a.) 6 mg/L Rifampicin, b.) 8 mg/L Rifampicin, c.) 12 mg/L Rifampicin. Post-adjustment model outputs for susceptible and resistant populations of E. coli grown in media containing d.) 6 mg/L Rifampicin, e.) 8 mg/L Rifampicin, f.) 12 mg/L Rifampicin.

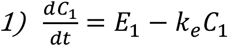

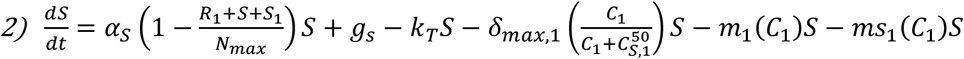

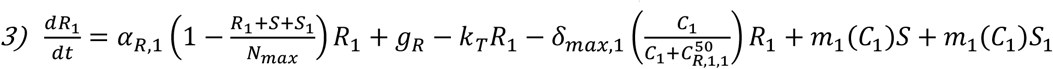

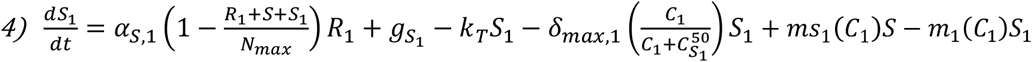

**Eq Set 2**. Augmented model of sensitive, semi-resistant, and resistant populations under selective pressure from Rifampicin therapy including only chromosomal mutation mechanisms

After adjustments, the resulting model outputs matched the general behavior of the resistant and susceptible bacterial populations in response to the selected concentrations of Rifampicin (Figure 2d-f). This established the ability of the model to match in vitro behavior allowing for the development of a model with predictive capabilities.

Following this initial study, the adjusted model was used to make predictions for a second experiment with different Rifampicin concentrations (Figure 3a-b). This second experiment was again conducted using the eVOLVER with the same growth conditions and the results of the experiment qualitatively matched the behavior predicted by the model (Figure 3c-d). This predictive capability demonstrated the validity of the modelling assumptions and prompted further simulation studies using the model.

**Figure 3.**
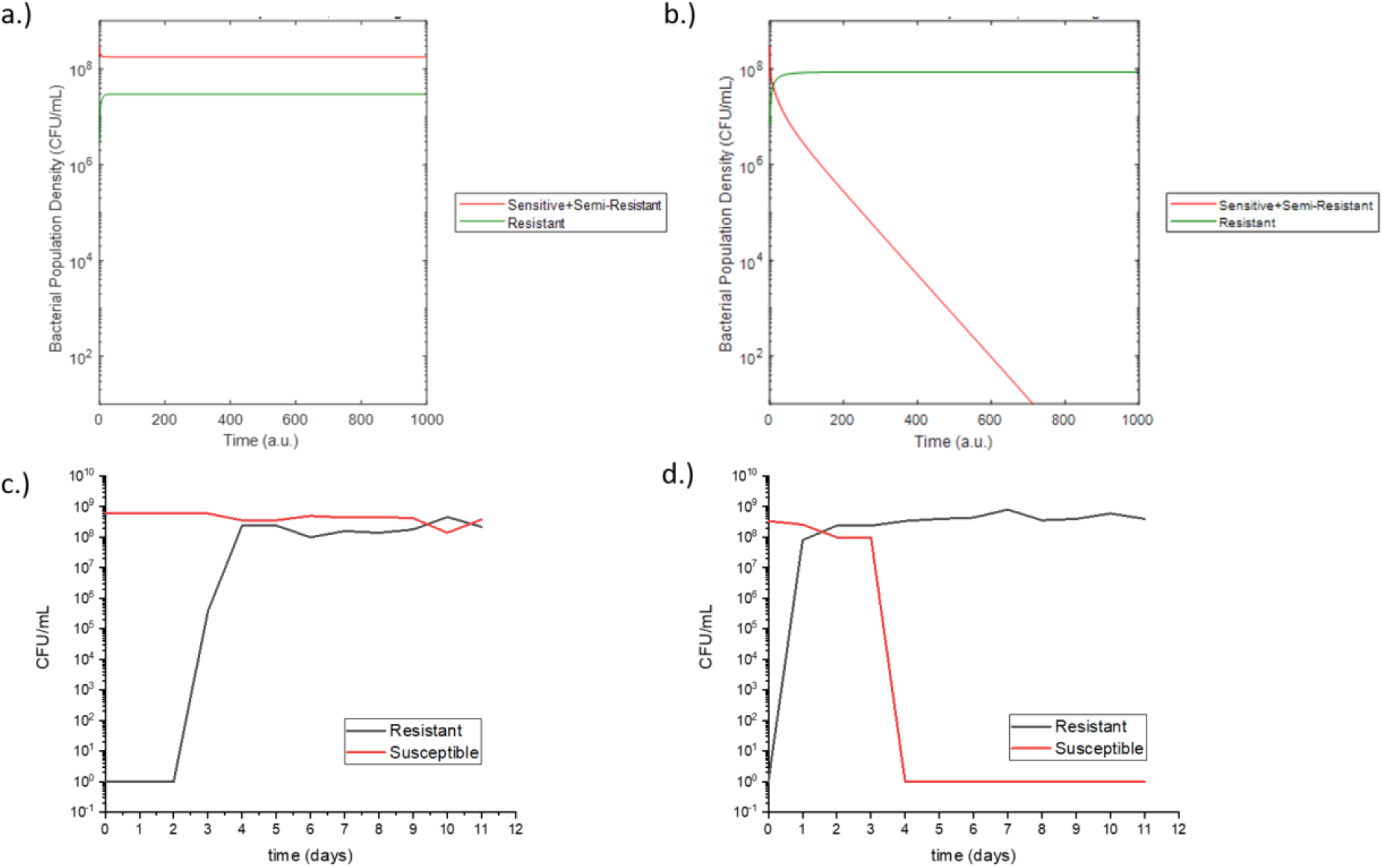
Post-adjustment model predictions for susceptible and resistant populations of *E. coli* grown in media containing a.) 3 mg/L Rifampicin, b.) 10 mg/L Rifampicin. Measured susceptible and resistant populations from eVOLVER experiments of E. coli grown in continuous culture in media containing c.) 3 mg/L Rifampicin, d.) 10 mg/L Rifampicin.

### Model Simulations

Initial model simulations were done to compare the outputs of our model with known scenarios under low antibiotic pressure. In the scenario of no HGT, as expected, only mutational resistance was observed (Figure 4a). Likewise, for a scenario in which there is no chromosomal mutation (Chromosomal Mutation Rate = Initial Mutant Population = 0), only resistance from HGT was observed and due to an assumed higher growth rate due to a lower fitness cost for bacteria with resistance-conferring plasmids than those with chromosomal resistance mutations (26), this resistance overtook the susceptible population at a higher rate (Figure 4b).

**Figure 4.**
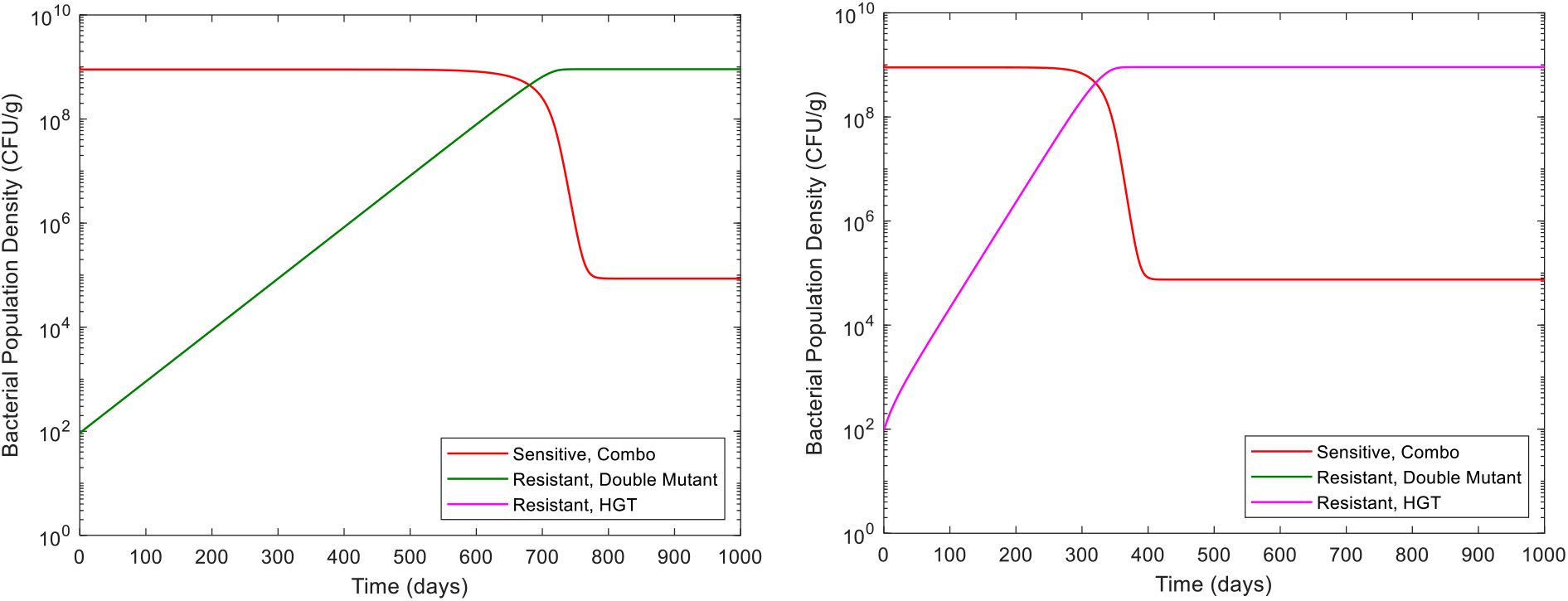
Sensitive and resistant populations under **a.)** resistance only from chromosomal mutations; **b.)** resistance only from the HGT of resistance conferring plasmids (MDR plasmid)

### Very low antibiotic residue concentration can promote resistant population growth

We then turned our attention to simulating realistic scenarios in wastewater and sewage. We studied the effects of changing antibiotic residue concentration on the development of antibiotic resistance (Figure 5). Low subinhibitory concentrations (0.5 - 2 ug/ml), as would be present in wastewater settings (4, 5), were studied. These concentrations are an order of magnitude lower than the minimum inhibitory concentrations of each of the two simulated antibiotics, which have been assumed as 25 ug/mL based on our experimental results with rifampicin. At these low concentrations, increased antibiotic residue concentration increased the rate of resistance development and decreased the time to resistant population dominance (defined as time when *R*_*m*_*+R*_*p*_*> S*). This is in agreement with several prior studies linking subtherapeutic antibiotic levels with both chromosomal resistance mutation development and HGT (6, 7, 9-12). Future studies will look at higher antibiotic concentrations and the point of inflection where this increase in antibiotic concentration begins to prevent resistance through bactericidal activity. Furthermore, HGT was observed to be the dominant mode of resistance, in agreement with patterns reported in studies of *E. coli* in other settings (20).

**Figure 5.**
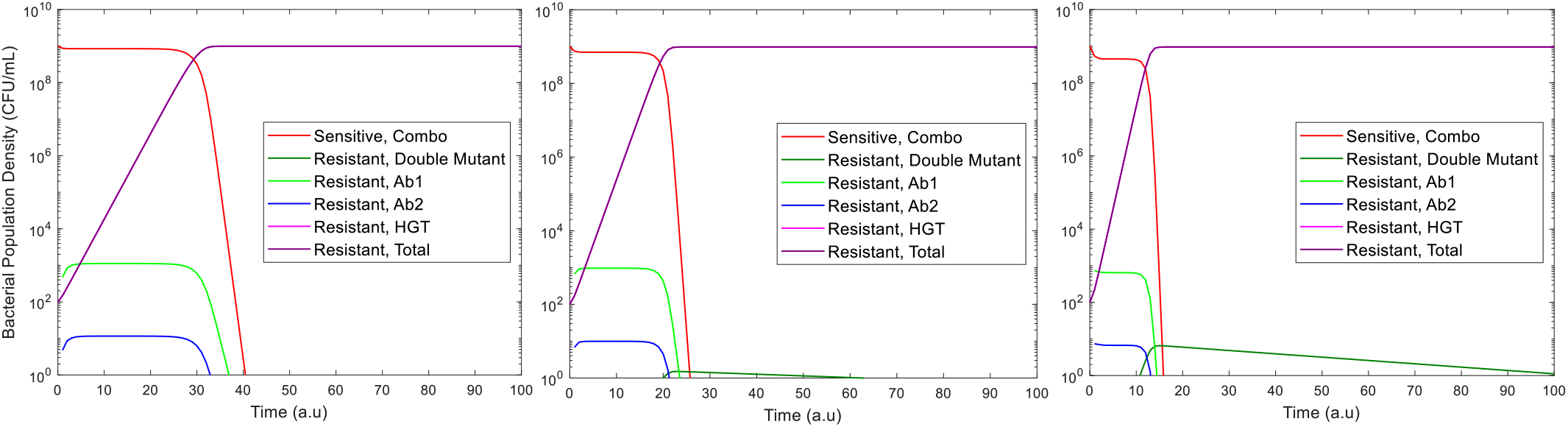
Sensitive and resistant populations under antibiotic residue concentrations of a.) 0.5 ug/ml antibiotic 1 + 0.5 ug/ml antibiotic 2; b.) 1 ug/ml antibiotic 1 + 1 ug/ml antibiotic 2; c.) 2 ug/ml antibiotic 1 + 2 ug/ml antibiotic 2

### Increase in HGT Rate increases resistance at low concentrations

Additionally, the effect of HGT was modelled by increasing the HGT rate, β (Figure 6). Prior studies have indicated that HGT rate is a less significant driver of resistance frequencies (21). However, our model shows that at very low subinhibitory concentrations of antibiotic, increased HGT rate significantly decreases the time to resistant population domination. This result indicates that increasing HGT rate allows resistance to be acquired at very low selective pressures where there are infrequent chromosomal mutations. This result is significant as it shows that decreasing antibiotic levels in wastewater to low levels is not sufficient in preventing the growth of resistant populations. At very low concentrations of antibiotics, the HGT rate of antibiotic resistance genes in wastewater can significantly affect the proliferation of resistant populations in bacteria with high gene transfer rates. As the HGT rates of the multitude of bacterial species in wastewater is not a commonly monitored parameter, this may be a factor to be mindful of in wastewater surveillance, as these rates can have a significant effect on the evolution of resistant population reservoirs even at low antibiotic concentrations.

**Figure 6.**
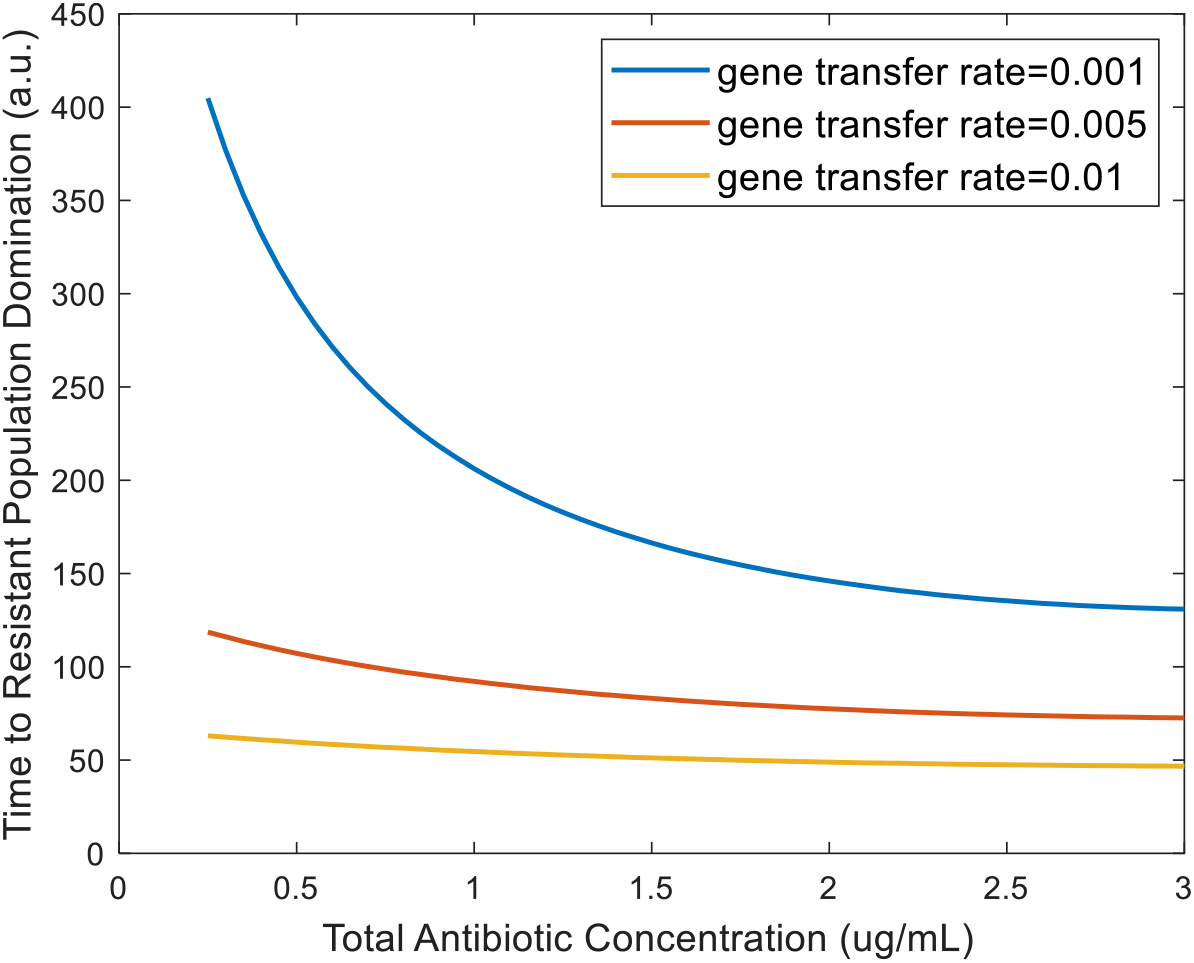
Effect of horizontal gene transfer rate on time to resistant population dominance for subinhibitory antibiotic residue concentrations (defined as time when *R*_*m*_+*R*_*p*_> *S*)

### Effect of Bacterial Killing Rate

Because wastewater settings can often have low concentrations of antibiotics from various polluting factors (4, 5), we probed the specific effects of antibiotic residues on the development of resistant populations. Individual antibiotics have different killing rates based on factors such as their modes of action. Thus, we investigated the effect of bacterial killing rate on resistant population growth (Figure 7). Increasing the killing rate terms (*δ*_*max*,1_* and δ*_*max*,2_) was observed to decrease the time it takes for the resistant population to dominate under subinhibitory concentrations. This result is in agreement with previous studies showing increased selective pressure from antibiotics at subinhibitory concentrations can increase resistance development (6, 7).

**Figure 7.**
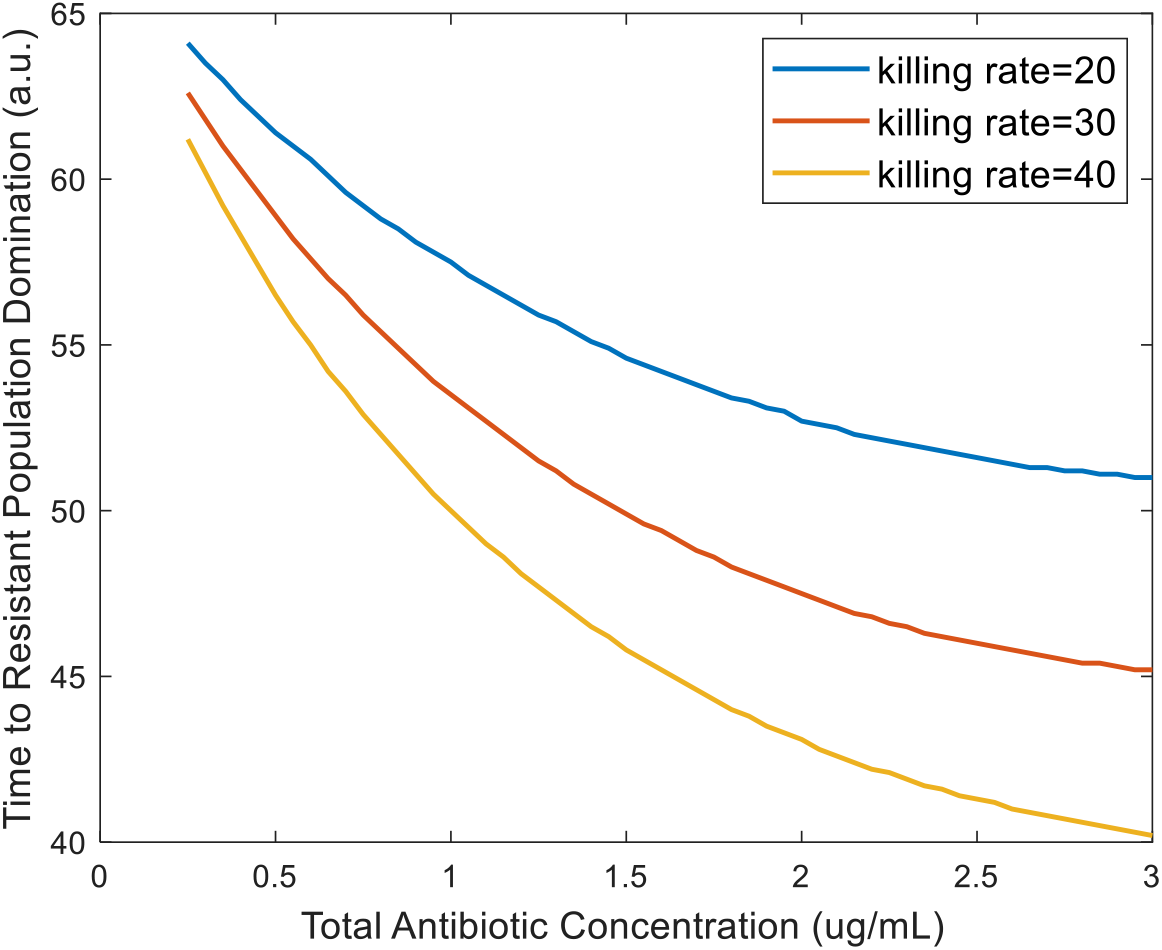
Effect of bacterial killing rate on time to resistant population dominance for subinhibitory antibiotic residue concentrations (defined as time when *R*_*m*_+*R*_*p*_> *S*)

### Synergistic antibiotic interactions increase resistant population growth

The interaction between two antibiotics has previously been shown to affect resistance acquisition (22). Synergy is the interaction of multiple drugs to have a greater killing action than the sum of their parts. Antibiotic synergy has also been shown to increase the likelihood of resistance acquisition at subtherapeutic doses (8). However, the effects of antibiotic interaction on the growth of resistant populations in wastewater settings, where many antibiotic residues can be present, has not previously been observed or modeled. In order to fill this gap, we have probed the effects of synergistic effects between antibiotic residues on the system by varying the synergy parameter, *syn*. Results from our model show that increased synergy between different antibiotics resulted in a decrease in the time to resistant population elimination at suprainhibitory concentrations (Figure 8a). This result may initially seem counterintuitive, but is in fact consistent with previous studies. This is because the synergistic action between the antibiotics increases the killing action to the point where it is effective on the resistant populations at lower concentrations than for antagonistic pairs. Hence, the synergy between the two antibiotics allows for greater bactericidal activity at lower concentrations. However, at subinhibitory concentrations, increasing synergy between antibiotics decreased the time to resistant population dominance (Figure 8b). This is in agreement with other models in which synergy between antibiotics was observed to increase resistance acquisition (8). Our model further shows that these effects of synergy are observable even with low levels of antibiotic, such as the antibiotic residue concentrations present in wastewater. While the effects of antibiotic interaction on resistance acquisition have been previously modeled, low subinhibitory antibiotic concentrations such as those found in wastewater have not been well studied and are critical for understanding resistance development in wastewater settings (17).

**Figure 8.**
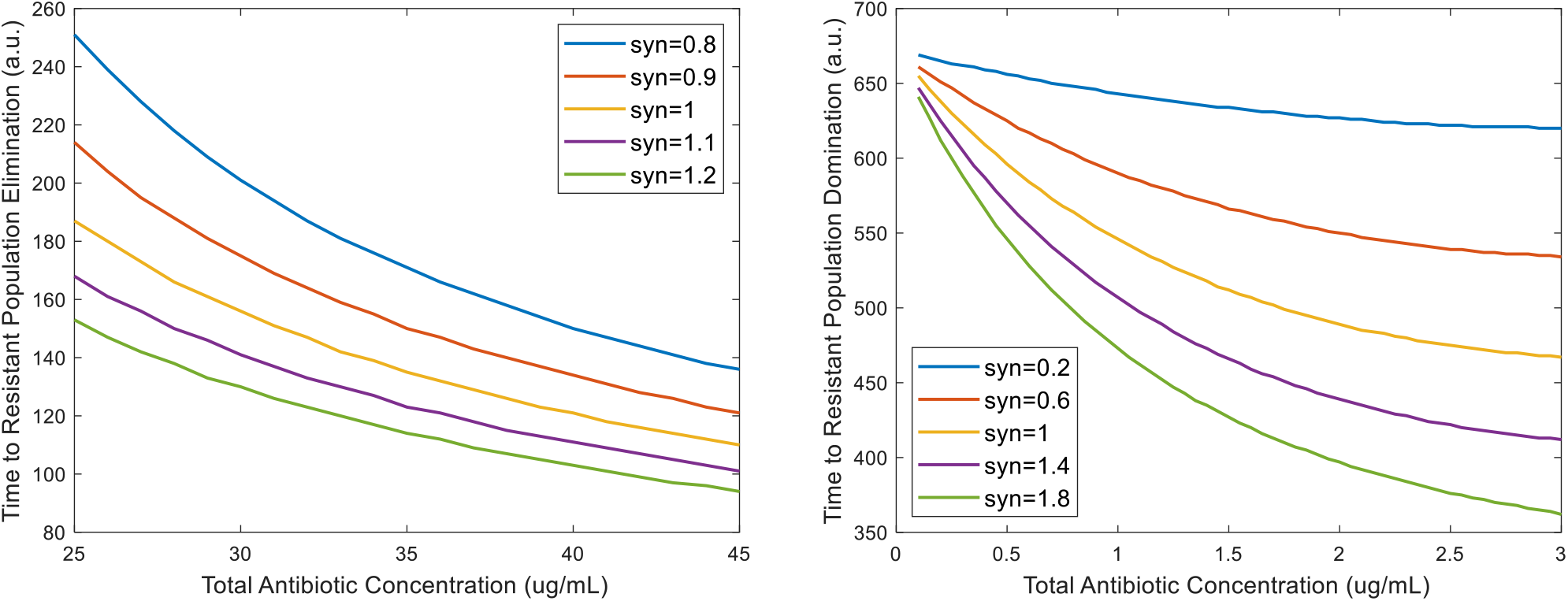
Effect of synergy of antibiotic combinations on **a.)** time to resistant population elimination for suprainhibitory antibiotic residue concentrations **b.)** time to resistant population dominance for subinhibitory antibiotic residue concentrations (defined as time when *R*_*m*_+*R*_p_> *S*)

## Conclusion

Though we have been able to draw a number of conclusions on the relative effect of a variety of factors affecting resistance development in wastewater, we note that our model does have limitations. First, our model is based on parameter values from literature, not all of which have been experimentally validated against wastewater conditions. Additionally, other phenomena related to microbial growth, such as dormancy, biofilm formation and the Long-Term Survival phase, have not been included in this initial model and the addition of these terms in future modelling studies may allow for better quantification of resistance development. However, despite these limitations, experimental validation demonstrated the ability of our model to qualitatively predict *in vitro* behavior of bacterial resistance development in response to multiple subinhibitory concentrations of Rifampicin (Figure 3). We have also assumed a simplified case where the two mechanisms of resistance acquisition that are modelled are independent, that as interaction between HGT and chromosomal mutation and the behavior of cells having both are unknown. This assumption can be tested through future experimental work. Our model also does not account for multiple bacterial species or a multitude of antibiotic residues, as would be present in a complex environment like wastewater. Interaction between the multiple bacterial species as well as quorum sensing within bacteria of the same species could have an effect on resistant population size and is a future area of expansion for this model. Additional improvements to the accuracy and robustness of the model could be made through parameter initialization from field data, and experimental validation of model inputs and outputs under specific wastewater conditions.

Despite these limitations, our model provides new, and important quantitative insight on the evolution of resistant bacterial reservoirs. It also provides an integrated framework to incorporate several aspects of resistance acquisition and growth previously lacking from models focused on AMR in wastewater settings. In terms of results, we have shown that HGT rate can be a significant driver of resistant population growth at very low antibiotic concentrations (Figures 5 and 6). This indicates that HGT rates of bacteria in wastewater may be important to monitor in addition to antibiotic residue concentration. We have also shown that synergy between the antibiotic residues present in wastewater can increase the rate of resistant population growth, even at the low concentrations present in wastewater (Figure 8). Thus, antibiotic residues in wastewater may pose a greater risk than might be expected without taking these antibiotic interactions into consideration. This has important implications for determining acceptable antibiotic levels in wastewater post treatment, as determining levels without considering antibiotic interactions may lead to overestimating the permissible level of antibiotic and allow for the proliferation of antibiotic resistant bacterial populations. This model can be adapted for use as a prediction tool for public health policy makers and be used to predict resistant population emergence in different sewage and wastewater conditions. Additionally, it can be expanded to be used to model different resistant outbreak prevention strategies in sewage and wastewater treatment.

## Author Contributions

I. Sutradhar designed the model, conducted experiments, and analyzed the data. C. Ching andD. Desai provided guidance on model design and verification. M. Suprenant provided guidance on data analysis. A. Khalil provided facilities for experimental work with the eVOLVER and E. Briars and Z. Heins provided assistance on experimental design. I. Sutradhar and M. H. Zaman wrote the article.

## Acknowledgements

This research was supported in part by the National Institutes of Health training grant at Boston University, T32 EB006359. A.S.K. also acknowledges funding from the NIH Director’s New Innovator Award (1DP2AI131083). The content is solely the responsibility of the authors and does not necessarily represent the official views of the National Institutes of Health.

## Notes

### Competing Interest Statement

The authors have declared no competing interest.

### Summary of Updates

Section added on experimental validation of the computational model; authors updated.

